# First isolation and genetic characterization of Puumala orthohantavirus strains from France

**DOI:** 10.1101/2020.08.27.270181

**Authors:** Johann Vulin, Séverine Murri, Sarah Madrières, Maxime Galan, Caroline Tatard, Sylvain Piry, Gabriele Vaccari, Claudia D’agostino, Nathalie Charbonnel, Guillaume Castel, Philippe Marianneau

**Affiliations:** Laboratoire de Lyon, ANSES, Unité de virologie, Lyon, France; CBGP, INRAE, CIRAD, IRD, Institut Agro, Univ Montpellier, Montpellier, France; Department of Food Safety, Nutrition and Veterinary Public Health, Istituto Superiore di Sanità, 00161 Rome, Italy

**Keywords:** Hantavirus, Puumala, isolation

## Abstract

Puumala orthohantavirus (PUUV) causes a mild form of haemorrhagic fever with renal syndrome (HFRS) called nephropathia epidemica (NE), regularly diagnosed in Europe. France represents the western frontier of the expansion of NE in Europe with two distinct areas: an endemic area (north-eastern France) where PUUV circulates in rodent populations, with detection of many human NE cases, and a non-endemic area (south-western France) where the virus is not detected, with only a few human cases being reported. France is thus a relevant country in which to study the factors that influence the evolution of PUUV distribution. In this study, we describe for the first time the isolation of two PUUV strains from two distinct French geographical areas: Ardennes (endemic area) and Loiret (non-endemic area). To isolate PUUV efficiently, we selected wild bank voles (*Myodes glareolus*, the specific reservoir of PUUV) captured in these areas and that were seronegative for anti-PUUV IgG (ELISA), but showed a non-negligible viral RNA load in their lung tissue (qRT-PCR). With this study design, we were able to cultivate and maintain these two strains in Vero E6 cells and also propagate both strains in immunologically neutral bank voles efficiently and rapidly. Complete coding sequences of the S and M segments were determined by Sanger sequencing from RNA extracted from positive bank voles (naturally and experimentally infected) and from supernatants of Vero E6 cell extracts. For the M segment, nucleotide sequences were 100% identical for both strains. For the S segment, the amino-acid sequences from each strain revealed one mismatch between sequences obtained from tissue and from cell supernatants, revealing distinct “bank vole” and a “cell” molecular profile. High-throughput sequencing confirmed Sanger results, and provided a better assessment of the impact of isolation methods on intra-host viral diversity.

## Text

Orthohantaviruses represent an increasing threat to humans due to their worldwide distribution, the increase in the number of infections and the emergence or re-emergence of new viruses (1). Puumala virus (PUUV) is the main orthohantavirus circulating in France. This tri-segmented, enveloped RNA virus, hosted by a rodent, the bank vole (*Myodes glaerolus*), causes a mild form of haemorrhagic fever with renal syndrome (HFRS) called nephropathia epidemica (NE) (2). About 100 cases of NE are reported annually in north-eastern France (3).

In France, PUUV sequences have been studied in bank vole samples (4) and more recently in human samples (5), but isolated strains have never been cultured or maintained until now. Considering the adaptation capacity of PUUV to cell culture conditions and changes in infectivity induced by cell culture passages (6), obtaining a “wild-type” virus is a crucial step towards a better understanding of the biology of this virus. Unfortunately, due to its slow growth in cell culture, PUUV is often very difficult to isolate (7).

In this study, we set out to determine a method for selecting bank voles containing live virus from which to isolate PUUV strains. Seto et al. (8) isolated PUUV from bank voles that harboured the orthohantavirus nucleocapsid protein (NP) in their lungs, but showed no antibodies against PUUV. This detection of infection coupled with the absence of antibodies probably corresponds to an active viraemic phase. We adapted the Seto et al. approach by screening our collections of bank vole samples (composed of voles captured in various French forests over the last 10 years (4) (9)) for seronegative animals with high amounts of viral RNA in their lung tissue. Serum samples were assayed using IgG ELISA as already described (9) and viral RNA was extracted from lung homogenates using the QIAamp Viral Mini Kit according to the manufacturer’s instructions (Qiagen). Quantitative RT-PCR was performed with 2.5 μL of viral RNA amplified using the SuperScript III One-Step RT-PCR system with Platinum Taq High Fidelity (Invitrogen) on a LightCycler 480 (Roche).

More specifically, we screened bank voles captured in two distinct French geographical areas: (1) an NE endemic area: Ardennes, where PUUV circulates in rodent populations and where many cases of human NE are detected (3); (2) a non-endemic zone: Loiret, where PUUV circulates in rodent populations and where no or very few cases of NE are diagnosed (4). We selected seven bank voles captured in Ardennes during autumn 2011 (out of a total of 201 animals captured during this trapping session) and one bank vole captured in Loiret in summer 2014 (out of 44 animals). For our first isolation assays, we used a sample from Ardennes named Hargnies/2011 with the smallest cycle threshold value (Ct; i.e. 18.90 cycles) and a lung tissue sample from Loiret named Vouzon/2014 that showed a Ct of 23.65 cycles.

From these two animals, we tried to isolate live viruses from lung homogenates (5% w/v). We tested many conditions, such as centrifugation (with or without), filtration (0.22 μM) and dilution (10^−1^ to 10^−4^) for homogenate preparation; cell confluence and numbers of cell passages before infection (16 or 43) for cell culture and different infection methods between the three passages before isolation (supernatant, transposition or co-culture). The conditions that led to the *in vitro* isolation of the two viral strains consisted of the use of the lung homogenate centrifuged (300 rpm for 10 min), not filtered, diluted to 10^−1^ and applied to low-passaged Vero E6 cells (16 passages) in DMEM (Dulbecco′s Modified Eagle′s – Medium) supplemented with 5% foetal calf serum (Gibco) and cultured to hyper-confluence (1×10^6^ cells for one well in P6).

It was necessary to carry out three successive passages to reach a sufficient quantity of live viruses. These passages were carried out using “co-culture”: a quarter of the initially infected cells were transferred to a culture of non-infected cells (2×10^5^ cells/well in P6) accompanied by 500 μL of the supernatant from the infected culture added to 2 mL of fresh media.

The viral load measured in the supernatant of Vero E6 cells increased substantially: −15.2 Ct and −12.8 Ct for Ardennes - Hargnies and Loiret – Vouzon, respectively **(Figure 1A)**. Moreover, at the end of cell isolation, viral titre was 1.6×10^4^ focus forming units (FFU)/mL and 1.5×10^4^ PFU/mL for Ardennes - Hargnies and Loiret - Vouzon, respectively. After titration, their focus sizes were different to that of the PUUV control (Sotkamo strain) serially passaged in Vero E6 cells **(Figure 1B)**. The focus size of cultivated viruses was smaller (in particular for Loiret - Vouzon) than Sotkamo foci. Phenotypic variations have already been described for PUUV titration assays (10). Altogether, these results show that we were able to select viral isolates from natural host tissue and cultivate them in cell culture.

**Figure 1:**
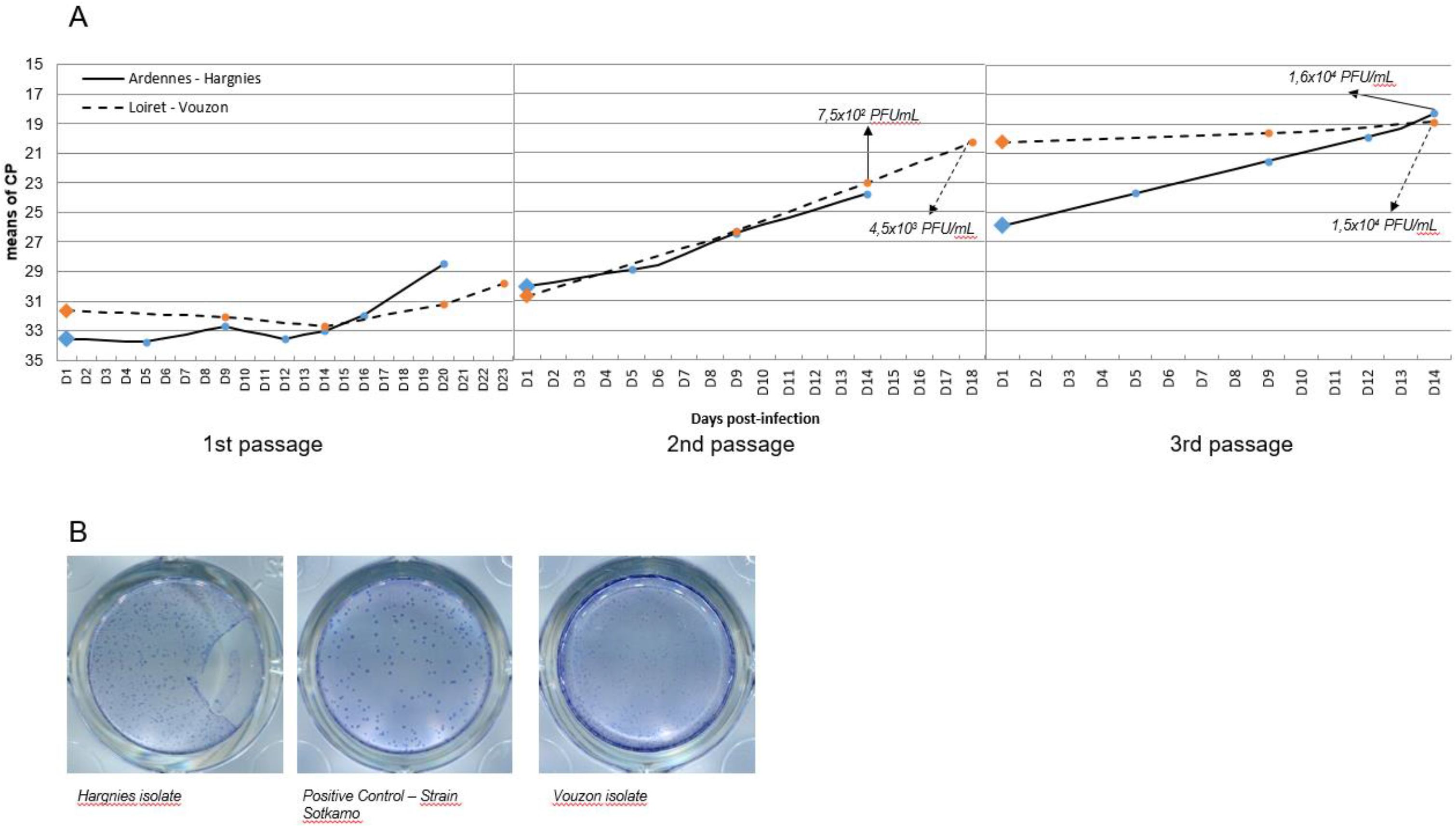
PUUV isolation with Vero E6 cells. Kinetics of PUUV RNA quantity during isolation process (**A**). Circles represent supernatants sampling. Picture of viral titration results (**B**) for Hargnies and Vouzon compared to Sotkamo referential strain.

Furthermore, we carried out an *in vivo* isolation process that consists of biological assays on bank voles that were maintained at the ISS Institute (Rome, Italy). We used the same lung homogenates already used for the *in vitro* isolation process. We performed a subcutaneous injection of lung homogenate (5% w/v) on two animals for each strain. None of the inoculated animals showed clinical evidence of viral infection. Seven days after infection, we collected blood samples via retro-orbital sinus puncture and carried out PUUV serological detection and viral RNA analyses. We did not detect any anti-PUUV antibodies (**Table 1**), but we found viral RNA in all sera analysed. Viral loads were similar regardless of the PUUV strain (Ardennes - Hargnies and Loiret - Vouzon), indicating that our protocol led to the effective transmission of PUUV in these animals. Two days later, at nine days post-infection (D9), we euthanized the four bank voles by cervical dislocation. We found that all animals had seroconverted, with a higher antibody titre observed for the Loiret - Vouzon strain (1/800) than for the Ardennes - Hargnies strain (animal 1: 1/200 and animal 2: 1/100). The RNA viral loads in sera had already decreased in all four animals analysed, but were still positive. We also analysed RNA viral loads in the lungs and liver, *i.e.* organs in which PUUV antigens have previously been detected (11, 12). We found high viral RNA loads (**Table 2**), which confirmed the effective infection of all animals by PUUV.

**Table 1:** Serological results. Serological results of bank vole experimentally infected with lung homogenates from animals trapped from Hargnies and Vouzon. OD: Optical Density – ND: Not Defined

| Capture area | Day post-infection | Animal number | Antibody IgG-NPuu |  |  |
| --- | --- | --- | --- | --- | --- |
|  |  |  | IgG-ELISA |  | Titration |
|  |  |  | OD | Conclusion | Last + dilution (OD) |
| Ardennes - Hagnies | D7 | 811 | 0.09 | - | ND |
|  | D7 | 812 | 0.058 | - | ND |
|  | D9 | 811 | 0.14 | + | 200 (0.102) |
|  | D9 | 812 | 0.117 | + | 100 (0.104) |
| Loiret - Vouzon | D7 | 791 | 0.032 | - | ND |
|  | D7 | 792 | 0.029 | - | ND |
|  | D9 | 791 | 0.23 | + | 800 (0.105) |
|  | D9 | 792 | 0.227 | + | 800 (0.106) |

**Table 2:** RNA detection results. RNA detection results of bank vole experimentally infected with lung homogenates from trapped animals at Hargnies and Vouzon.

| Strain | Day post-infection | Animal number | RT-PCR (mean Ct) |  |  |  |
| --- | --- | --- | --- | --- | --- | --- |
|  |  |  | Lung | Liver | Sera - Day7 | Sera - Day 9 |
| Ardennes - Hargnies | D9 | 811 | 16.5 | 19.7 | 29.2 | 32.1 |
|  |  | 812 | 19.1 | 20,0 | 30.6 | 32.7 |
| Loiret - Vouzon | D9 | 791 | 19.0 | 21,0 | 30.7 | 31,0 |
|  |  | 792 | 19.1 | 20.8 | 29.9 | 30.5 |

Molecular analyses were carried out as already described (7) to assess PUUV levels during bank vole infection. RNA was extracted from lung tissue samples of positive bank voles (natural populations and experimentally infected) and from the supernatant of Vero E6 cells, then sequenced to obtain the complete coding sequence of the S and M segments (for a list of sequences and accession numbers, see Supplementary Table S1). RNA was amplified using RT-PCR and nested PCR as already described (4). PCR products were sequenced using the Sanger method. The S and M sequences from Ardennes - Hargnies and Loiret - Vouzon were aligned using the Clustal Omega alignment program implemented in Seaview 4.5.0 (13). Nucleotide sequences were translated into amino-acid sequences and analysed using SeaView 4.5.0. For the M segment (3444 bp for coding region), the PUUV nucleotide sequences from Ardennes - Hargnies and Loiret - Vouzon were 100% identical for sequences obtained from lung tissue or cell supernatant. For the S segment (1304 bp for coding region), the amino-acid sequences from Ardennes - Hargnies and Loiret - Vouzon revealed one mismatch between the sequences from lung tissue and cell supernatant (**Figure 2**). Therefore, isolation protocols (*in vitro* or *in vivo*) seemed to have influenced the amino-acid sequences of both PUUV strains.

**Figure 2:**
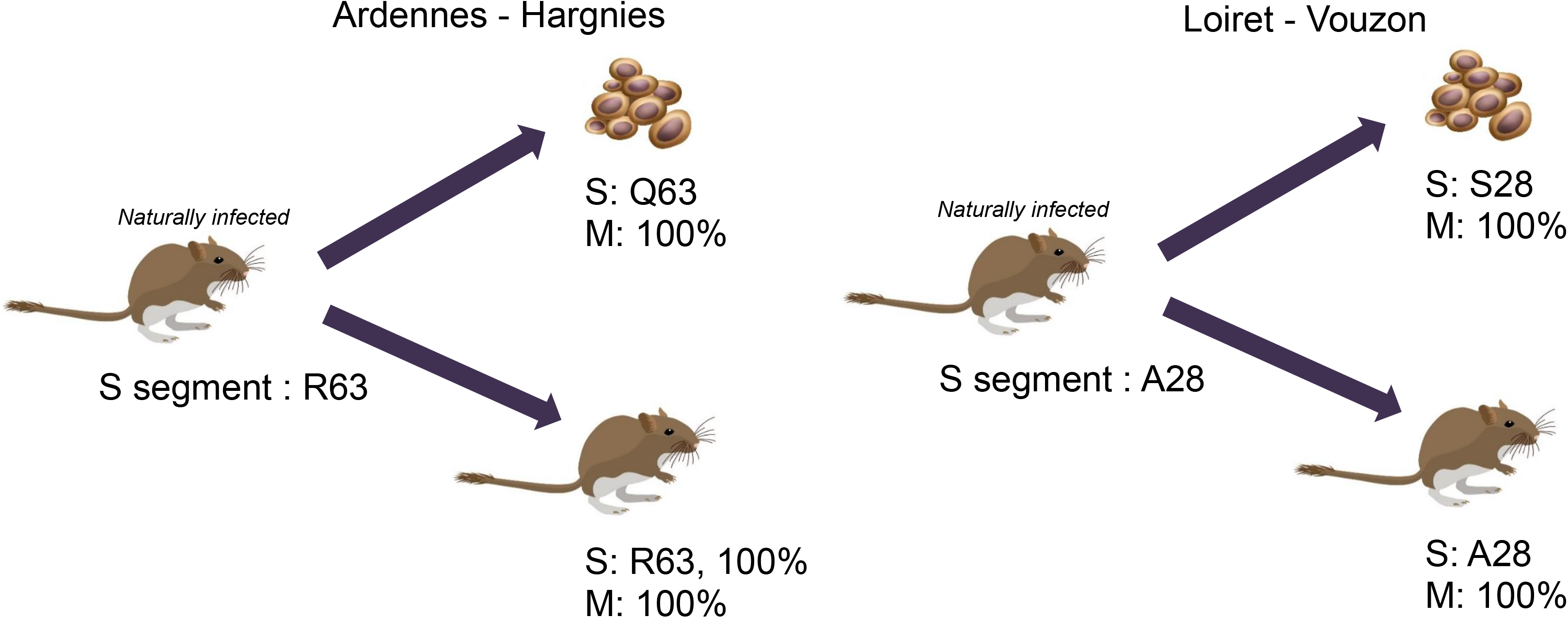
Comparison of S and M segment sequences. Synthesis of Sangers sequencing of S and M segments for Natural Isolates (sequence as reference), cell cultivated viruses and experimentally infected bank voles. Bank vole and cells drawing were designed by brgfx / Freepik

We then performed high-throughput sequencing (HTS) analyses to investigate these genetic variations further. More specifically, we assessed whether the different isolation protocols maintained the initial intra-host viral diversity detected in the original samples. We thus characterized the viral variants pool at each step of the isolation process. The PUUV S segment (1750 bp) was sequenced using MiSeq Illumina technology with 10 overlapping amplicons (named A to J) of about 250 bp (see Supplementary Table S2). To do so, 300 ng of viral RNA from supernatant of Vero E6 cells and 1500 ng from lung tissue of infected rodents were reverse-transcribed using the SuperScript III First-Strand Synthesis System (Invitrogen) according to the manufacturer’s instructions. The reverse-transcription reaction was performed with 2 μM of primers PUU1F1 (5’-CCTTGAAAAGCTACTACGAG-3’) and PUU1R1 (5’-CCTTGAAAAGCAATCAAGAA-3’) and with 50 ng of random hexamers provided in the kit. The sequencing libraries were prepared using a two-step PCR strategy adapted from Galan et al. (14) (see Supplementary information S3) and combined in a multiplex sequencing approach using unique dual indices (UDI) (15): each 9-bp i5 and i7 dual index was used only for one PCR sample without combinatorial indexing to ensure that libraries were sequenced and demultiplexed with the highest accuracy without any “leakage” between libraries (16). The libraries were sequenced on a MiSeq platform (GenSeq, Montpellier, France) with a 500-cycle reagent kit v2 (Illumina). A run of 250 bp paired-end sequencing was performed. The sequences were analysed with the data pre-processing tool FROGS (Genotoul) (17) and chimeric variants were removed using the isBimeraDenovo function of Dada2 R package (18). Each sample was analysed independently (qRT-PCRs, PCR amplifications and sequencing) using at least three PCR replicates to distinguish the true genetic variants and the artefactual mutations due to polymerase or sequencing errors. Mutations below a threshold of 0.32% were removed (see Supplementary information S4). Validated variants were aligned and analysed using SeaView 5.0 (13). Two indices were used to compare the viral diversity between the two isolation protocols and the two strains: the number of polymorphic sites (19, 20) and the mean percent complexity (the number of unique sequence reads/total reads × 100) calculated on the 10 amplicons (21). A Kruskal-Wallis test followed by a Dunn multiple comparison test was conducted to compare the mean percent complexity between strains of different origin (*in natura*, *in vitro* or *in vivo*).

*In natura*, the viral diversity of the Ardennes - Hargnies and Loiret - Vouzon strains were similar. We found no difference in the number of polymorphic sites or in the mean percent complexity between areas. After the *in vitro* isolation process, one variant of the Ardennes - Hargnies strain (R63), mostly found *in natura*, switch to another major variant, such as Q63. Nevertheless, R63 was still present at a high frequency (about 23%) instead of a Viral diversity decreased after the cell culture passages, regardless of the diversity index considered. The Loiret - Vouzon strain, as observed with Sanger method, showed a change in the major variant from A28 to S28. However, HTS results were contradictory: the number of polymorphic sites increased, but the mean percent complexity remained stable. The *in vitro* isolation process lead to a significant difference in viral diversity between PUUV strains, but not in the same way for both strains **(Figure 3A)**.

**Figure 3:**
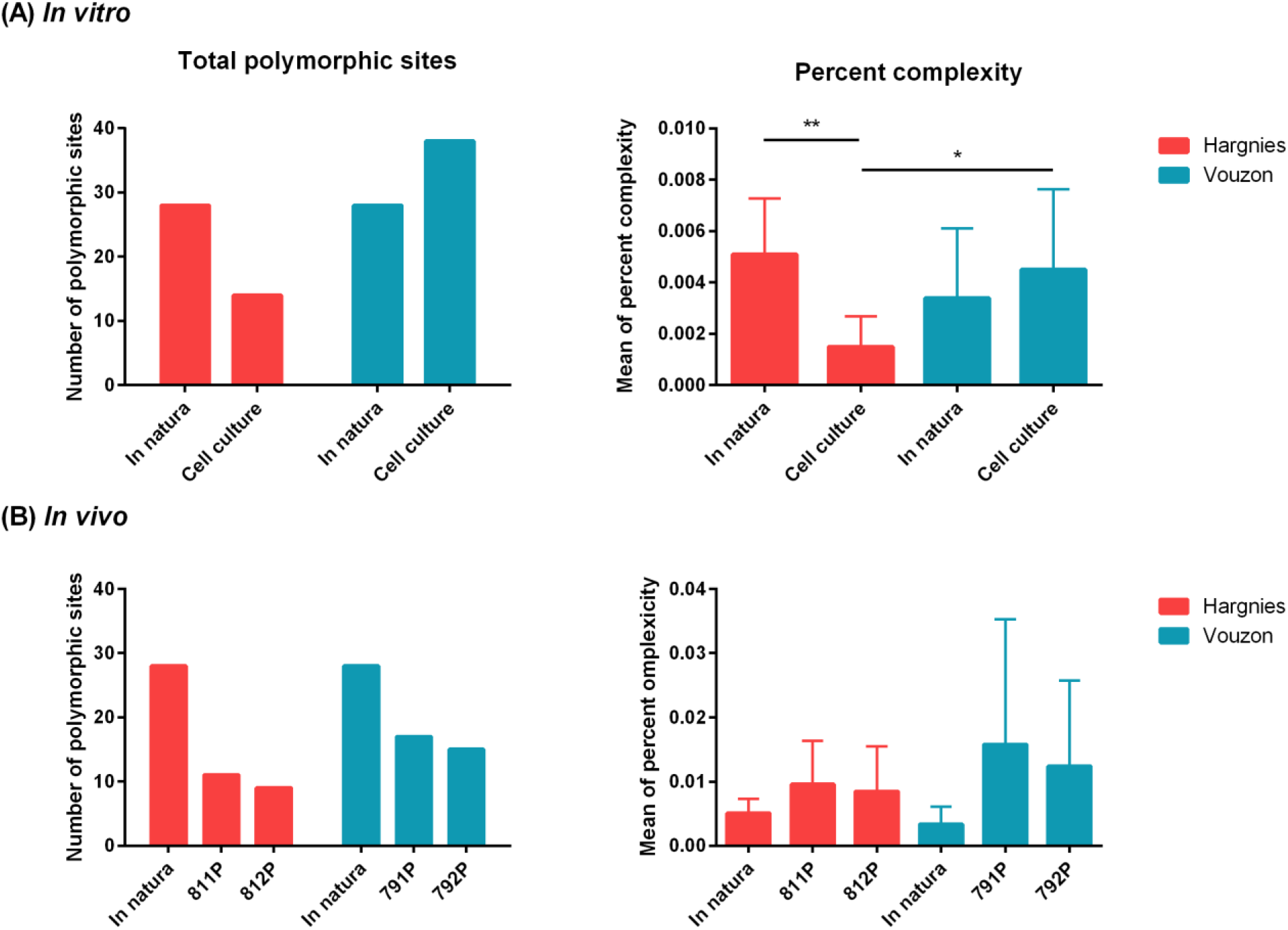
Comparison of viral diversity after (**A**) *in vitro* and (**B**) *in vivo* isolation process.

After the *in vivo* isolation process, we detected a lower number of polymorphic sites compared with the natural strains. However, we found no significant difference in the mean percent complexity between PUUV strains or in *in natura* and *in vitro* conditions **(Figure 3B)**. The sequence of the majority variant did not differ between these two conditions, regardless of the isolation protocol considered. Results were similar in bank voles infected with the same strain.

## Conclusion

In this study, we were able, for the first time, to cultivate and maintain in cell culture two PUUV isolates from two distinct French areas. Molecular analyses of the S and M segments of PUUV originating from natural isolates, from experimentally infected bank voles and from cell cultures revealed only one amino-acid mismatch for the Ardennes – Hargnies and Loiret - Vouzon strains. Both mismatches were identified in cell culture. HTS confirmed this result, thereby leading to the definition of a “bank vole” and a “cell culture” molecular profile. Having these two types of PUUV wild strain cultures is an important asset. Cell-adapted strains provide well-characterized viruses that can be used for antiviral candidate studies or cell-level experiments to assess the role of apoptosis, PUUV propagation or control (22). Furthermore, wild strains maintained in their natural host can help contribute to improving our knowledge of PUUV ecology and evolution (23). The bank voles used in the *in vivo* experiments came from a “neutral” origin, *i.e.* not from Ardennes or Loiret. For the moment, bank vole-PUUV interactions are poorly documented, but may be a key point in viral diversity.

This study also provides new evidence for a better understanding of regional epidemiological differences regarding the circulation of these PUUV strains in humans and in bank voles (4). Previous studies showed that there is significant inter-regional viral genetic diversity (4, 5, 24), which may explain, at least in part, the regional differences in NE epidemiological status in France. Here, we showed that the viral diversity of PUUV circulating in the NE endemic area (Ardennes) and in the NE non-endemic area (Loiret) evolve differently when passaged on Vero E6 cells. Further research dedicated to the characterization of the now-available French Puumala strains will help further our knowledge on the NE epidemiological situation.

## Ethics statement

All animal studies were conducted in accordance with French and European regulations on the care and protection of laboratory animals (French Decree No 2013-118 of 1 February 2013 and Directive 2010/63/EU of 22 September 2010).

## Acknowledgments

Data used in this work were partly produced through sequencing facilities at ISEM (Institut des Sciences et de l’Evolution-Montpellier) and the LabEx CEMEB (Centre Méditérranéen de l’Environnement et de la Biodiversité). Sarah Madrières was funded by an INRA-EFPA/ANSES fellowship.

