## supplementary information for "First isolation and genetic characterization of Puumala orthohantavirus strains from France"

### Supplementary material

#### Supplementary Table S1: PUUV sequences obtained in this study

| Strain or isolate[†] name | GenBank Accession No (S segment) | GenBank Accession No (M segment) | Sampling Year | GenBank Host |
|---|---|---|---|---|
| Hargnies-Mg | MW148463 | MW148469 | 2018 | *Myodes_glareolus* |
| Vouzon-Mg | MW148464 | MW148470 | 2018 | *Myodes_glareolus* |
| Hargnies-E6 | MW148465 | MW148471 | 2017 | Vero_E6_cells |
| Ard161_Ardennes_Mg_2011[†] | MW148466 | MW148472 | 2011 | *Myodes_glareolus* |
| Vouzon-E6 | MW148467 | MW148473 | 2017 | Vero_E6_cells |
| NCHA000357_Loiret_Mg_2014 [†] | MW148468 | MW148474 | 2014 | *Myodes_glareolus* |

#### Supplementary Table S2: Sequences of primer cocktails used in Miseq

| Amplicons | Primers names | Primer sequences (5' to 3') |
|---|---|---|
| A | PUMAMISEQ-Abis-F | GAGGATATAACCCGCCATGAA |
| | PUMAMISEQ-Abis1-F | GAGGTTATAACCCGCCATGAA |
| | PUMAMISEQ-Abis2-F | GAAGATATAACCCGCCATGAA |
| | PUMAMISEQ-A1-R | CAGTCGGGTCAGTAGGCTTA |
| | PUMAMISEQ-A2-R | CAGTCGGGTCAGCAGGCTTA |
| B | PUMAMISEQ-B1-F | TGGARGACAAACTTGCA |
| | PUMAMISEQ-B2-F | TGGARGATAAACTTGCA |
| | PUMAMISEQ-B1-R | ATRATAGGGAGTGTAAAGCC |
| | PUMAMISEQ-B2-R | ATRATAGGGAGTGTGAAGCC |
| | PUMAMISEQ-B3-R | ATRATAGGGAGTGTAAATCC |
| C | PUMAMISEQ-C1-F | GCCAAACAGCAGACTGGTA |
| | PUMAMISEQ-C2-F | GTCAAACAGCAGACTGGTA |
| | PUMAMISEQ-C3-F | GACAAACAGCAGATTGGTA |
| | PUMAMISEQ-C1-R | GTYARYTCYTCTGCTTTCATGGT |
| | PUMAMISEQ-C2-R | GTYARYTCYTCAGCCTTCATAGT |
| | PUMAMISEQ-C3-R | GTYARYTCYTCCGCCTTCATAGT |
| | PUMAMISEQ-C4-R | GTYARYTCYTCTGCCTTCATAGT |
| | PUMAMISEQ-C5-R | GTYARYTCYTCTGCTTTCATAGT |
| D | PUMAMISEQ-D1-F | GCATTAGAAGACCAAAGCACTT |
| | PUMAMISEQ-D2-F | GTATCAGGAGACCAAAGCATCT |
| | PUMAMISEQ-D3-F | GTATCCGAAGACCATAGCACTT |
| | PUMAMISEQ-D4-F | GTATTAGAAGACCAAAGCATTT |

| | | |
|---|---|---|
| | PUMAMISEQ-D5-F | GTATTCGCAGGCCAAAGCATCT |
| | PUMAMISEQ-D6-F | GTATTCGCAGGCCAAAGCACCT |
| | PUMAMISEQ-D1-R | GGTGTBCCCGGTTTTAT |
| | PUMAMISEQ-D2-R | GGTGTBCCTGGCTTCAC |
| | PUMAMISEQ-D3-R | GGTGTBCCGGGTTTCAC |
| | PUMAMISEQ-D4-R | GGTGTBCCCGGTTTCAT |
| | PUMAMISEQ-D5-R | GGTGTBCCTGGCTTGAC |
| | PUMAMISEQ-D6-R | GGTGTBCCTGGTTTGAC |
| | PUMAMISEQ-D7-R | GGTGTBCCTGGATTGAC |
| E | PUMAMISEQ-E1-F | TBAARGAYTGGGCAGACCGGA |
| | PUMAMISEQ-E2-F | TBAARGAYTGGGCAGATCGGA |
| | PUMAMISEQ-E3-F | TBAARGAYTGGACAGATCGGA |
| | PUMAMISEQ-E4-F | TBAARGAYTGGACAGACCGGA |
| | PUMAMISEQ-E1-R | CARGGTGCATTAGGAGAC |
| | PUMAMISEQ-E2-R | CARGGTGCATTTGGGGCT |
| | PUMAMISEQ-E3-R | CARGGTGCATTGGGTGAC |
| | PUMAMISEQ-E4-R | CARGGTGCATTAGGGGAC |
| F | PUMAMISEQ-F1-F | TYGAYTAYGCAGCCTCAGGA |
| | PUMAMISEQ-F2-F | TYGAYTAYGCAGCTTCCGGA |
| | PUMAMISEQ-F3-F | TYGAYTAYGCAGCCGCAGGA |
| | PUMAMISEQ-F1-R | GRGTYCKTCGCAGGTATG |
| | PUMAMISEQ-F2-R | GRGTYCKCCGCAGGTATG |
| | PUMAMISEQ-F3-R | GRGTYCKTCGCAAGTATG |
| | PUMAMISEQ-F4-R | GRGTYCKTCTTAGGTATG |
| G | PUMAMISEQ-G1-F | CAGGAYATGAGGAATACCATC |
| | PUMAMISEQ-G2-F | CAGGAYATGAGAAATACCATC |
| | PUMAMISEQ-G3-F | CAGGAYATGAGAAACACCATC |
| | PUMAMISEQ-G4-F | CAGGAYATGAGGAATACTATC |
| | PUMAMISEQ-G5-F | CAGGAYATGAGGAACACCATC |
| | PUMAMISEQ-G1-R | TYARGGGTTCCTGATTAGAA |
| | PUMAMISEQ-G2-R | TYARGGGCTCCTGGTTGGAG |
| | PUMAMISEQ-G3-R | TYARGGGCTCTTGATTGGAT |
| | PUMAMISEQ-G4-R | TYARGGGTTCCTGATTGGAA |
| | PUMAMISEQ-G5-R | TYARGGGCTCCTGGTTTGAT |
| | PUMAMISEQ-G6-R | TYARGGGCTCCTGCTTGGAG |
| H | PUMAMISEQ-H1-F | ATGGTRGAYCACTTTCATCTG |
| | PUMAMISEQ-H2-F | ATGGTRGAYCATTTCCACCTT |
| | PUMAMISEQ-H3-F | ATGGTRGAYCACTTTCACCTG |
| | PUMAMISEQ-H4-F | ATGGTRGAYCACTTTCATTTG |
| | PUMAMISEQ-H1-R | AGTTAAACCCTGATTAATCTAA |
| | PUMAMISEQ-H2-R | AGTTAAACCCTGATTAACCTGA |
| | PUMAMISEQ-H3-R | AGTTAAACCCTGATCAACCTAA |
| | PUMAMISEQ-H4-R | AGTTAAACCCTGATTGACCTTA |

| | | |
|---|---|---|
| I | PUMAMISEQ-I1-F | GGGATTAYYDTAATTAATTGTT |
| | PUMAMISEQ-I2-F | GGGATTAYYDTAGTTAATTGTT |
| | PUMAMISEQ-I3-F | GGGATTAYYDTAGTTAAGTGTT |
| | PUMAMISEQ-I4-F | GGGATTAYYDTAGTTGAATGTT |
| | PUMAMISEQ-I1-R | GCTCAGTTTCACATTATTGG |
| | PUMAMISEQ-I2-R | GCTTAGTTTCACATTATTGG |
| | PUMAMISEQ-I3-R | GCTCAGTTTCACATTAATGG |
| J | PUMAMISEQ-J1-F | TGCTGCTTAATCATTATTACCAGCA |
| | PUMAMISEQ-J2-F | TGCTGCTCAATTATTATCACCAGCT |
| | PUMAMISEQ-J3-F | TGCTATTGATAAGTGTTACCAGCA |
| | PUMAMISEQ-J4-F | TGCTGCTTAATCATTATACCCAGCA |
| | PUMAMISEQ-J5-F | TGCTGCTTGATTATTGTAACCAGCA |
| | PUMAMISEQ-J1-R | GAAATCAGTATGTTGAGGTAGT |
| | PUMAMISEQ-J2-R | GAAATCGGCATGTTGAGGTAGT |

**Supplementary information S3: Library preparation adapted from Galan et al. 2018**

PCR1 was performed in a 10 µL reaction volume using 5 µL of 2X Qiagen Multiplex Kit Master Mix (Qiagen), 0.3 µM of each primer and 2 µL of 4-fold diluted cDNA. PCR conditions consisted of an initial denaturation step at 95°C for 15 min, followed by 40 cycles of denaturation at 94°C for 30 s, annealing at 50°C for 1 min, and extension at 72°C for 40 s, followed by a final extension step at 72°C for 10 min.

The PCR2 consisted of a limited-cycle amplification step to add multiplexing indices i5 and i7 and Illumina sequencing adapters P5 and P7 at both ends of each cDNA fragment. PCR2 was carried out using 5 µL of 2X Qiagen Multiplex Kit Master Mix, 0.5 µM of each indexed primer and 2 µL of PCR1. Amplification started with an initial denaturation step of 95°C for 15 min, followed by 8 cycles of denaturation at 94°C for 40 s, annealing at 55°C for 1 min and extension at 72°C for 1 min, followed by a final extension step at 72°C for 10 min.

The average expected size of the super pool for excision on an agarose gel was 406 bp (including primers, indices and adaptors). xx 12 pM of library and 5% of PhiX control were loaded on a Miseq flow cell.

**Supplementary information S4: Definition of the 0.32% threshold**

Because plasmid DNA propagation has a much lower error rate (≈$10^{-7}$–$10^{-8}$ errors/site/copy) than RNA virus genome replication, a plasmid-generated “viral” RNA can be considered as an extremely low diversified starting population. This strategy can be used to determine the error rate and separate real mutations from process error (Orton et al., 2015).

In this study, the threshold of 0.32% was set after sequencing a plasmid generated from the Sotkamo PUUV strain. First, the complete PUUV S segment region (1759 bp) was amplified in 10 replicates with a final volume of 10 µL. We used 5 µL of 2X Qiagen Multiplex Kit Master Mix, 0.3µM of forward PUMAEXTLG-F (5'-CTGGAATGAGTGACTTGACAG-3') and reverse PUMAEXTLG-R (5'-CAGCATGTTGAGGTAGTATATTG-3') primers and 2 µL of 5-fold diluted Sotkamo cDNA. PCR conditions consisted in an initial denaturation step at 95°C for 15 min, followed by 40 cycles of denaturation at 94°C for 30 s, annealing at 45°C for 90 s, and extension at 72°C for 2 min, followed by a final extension step at 72°C for 10 min. Then, 5µL of PCR product was verified by electrophoresis on a 1.5% agarose gel. Then, all PCR products were pooled, purified with Ampure XP beads (Beckman coulter) and quantified with Qubit Fluorometer (ThermoFischer).

PCR products were cloned using the pGEM-T Easy vector system 2 (Promega) following the manufacturer's instructions. A 3:1 PCR product: vector molar ratio was used for ligation. Transformation was performed using JM109 High Efficiency competent cells growing on LB/Ampicillin/IPTG/X-Gal Petri plates overnight at 37°C. White bacterial colonies were picked and screened by PCR amplification using T7 and SP6 primers. Two recombinant plasmids were selected and purified using Qiaprep Spin Mini Prep (Qiagen) and then sequenced with Sanger technology at Eurofins MWG. Finally, 50-fold dilutions of the two purified plasmid were used as template for short amplification (Illumina) to define the cycle threshold.
